# Split-HaloTag^®^ Imaging Assay for Sophisticated Microscopy of Protein-Protein Interactions *in planta*

**DOI:** 10.1101/2020.06.08.139378

**Authors:** Rieke Minner-Meinen, Jan-Niklas Weber, Andreas Albrecht, Rainer Matis, Maria Behnecke, Cindy Tietge, Stefan Frank, Jutta Schulze, Henrik Buschmann, Peter Jomo Walla, Ralf-R. Mendel, Robert Hänsch, David Kaufholdt

## Abstract

An ever-increasing number of intracellular multi-protein networks have been identified in plant cells. Split-GFP based protein-protein interaction assays combine the advantages of *in vivo* interaction studies in a native environment with additional visualisation of protein complex localisation. Due to its simple protocols, it has become one of the most frequently used methods. However, standard fluorescent proteins entail several drawbacks for sophisticated microscopy.

With the HaloTag^®^ system, these drawbacks can be overcome as this reporter forms covalent irreversible bonds with synthetic photostable fluorescent ligands. Dyes can be used in adjustable concentrations and are suitable for advanced microscopy methods. Therefore, we established the Split-HaloTag^®^ imaging assay in plants which is based on the reconstitution of a functional HaloTag^®^ protein upon protein-protein interaction and subsequent covalent binding of an added fluorescent ligand. Its suitability and robustness were demonstrated using well-characterised interactions as an example for protein-protein interaction at cellular structures: the molybdenum cofactor biosynthesis complex anchoring to filamentous actin. Additionally, a specific interaction was visualised with subdiffractional polarisation microscopy in a more distinctive manner as example for sophisticated imaging.

Split-GFP and Split-HaloTag^®^ can complement one another as Split-HaloTag^®^ represents an alternative option and an addition to the large toolbox of *in vivo* methods. Therefore, this promising new Split-HaloTag^®^ imaging assay provides a unique and sensitive approach for more detailed characterization of protein-protein interaction with specific microscopic techniques such as 3D-imaging, single molecule tracking and super-resolution microscopy.

## Introduction

An ever-increasing number of protein networks has been identified in plants (Zitnik *et al*., 2019). Therefore, understanding the cellular biology of substrate channelling pathways requires the characterisation of protein-protein interactions (PPIs) in their native environment. A broad spectrum of *in vivo* methods has been employed to analyse PPIs such as bimolecular fluorescence complementation (BiFC) belonging to the group of protein fragment complementation assays (Struk *et al*., 2019). Basically, two non-fluorescent reporter fragments of a fluorescent protein (FP) are fused genetically to putative interaction partners and an interaction between the two allows formation of a bimolecular fluorescent complex (Kerppola, 2008). Consequently, BiFC not only allows detection of PPIs but also visualisation and localisation of the protein complex (Bhat *et al*., 2006). Furthermore, using FPs results in fluorescence signals without an invasive insertion of exogenous chemical compounds into the cell. Therefore, due to its simple protocols BiFC has become one of the most popular and frequently used method to study PPIs in plant cells (Kudla & Bock, 2016).

Conventional light and fluorescence microscopy are diffraction-limited to a resolution limit of approx. 200 nm in the lateral (x-y) and about 600 nm in the axial (z) direction (Cremer and Masters, 2013). Many subcellular structures are smaller which hampers their detailed observation (Huang *et al*., 2009). To circumvent these restrictions, advanced fluorescence imaging methods such as single molecule detection, subdiffractional polarisation imaging or super-resolution microscopy (SRM) techniques have been developed to improve the resolution and to allow studying molecular processes more detailed (Moerner and Kador 1989; Orrit and Bernard, 1990; Bode *et al*., 2008; Holleboom *et al*., 2014; Liao *et al*., 2010/2011; Godin *et al*., 2014; Loison *et al*., 2018; Camacho *et al*., 2019). However, for such advanced imaging techniques fluorescent dyes with high stability and brightness are needed (Banaz *et al*., 2018), which is hard to realise by standard FPs as they show low quantum efficiency, blinking behaviour, a high photobleaching rate during long-term observations (Reck-Petersen *et al*., 2006), photoswitching (Morisaki and McNally, 2014) as well as the tendency to form oligomers (Miyawaki *et al*., 2003).

Self-labelling enzyme tags such as the HaloTag^®^ have been shown to overcome these drawbacks of FPs and are suitable for such microscopy methods and super-resolution imaging (Grimm *et al*., 2015; Lee *et al*., 2010). The HaloTag^®^ System (Promega, https://www.promega.de/) is based on the bacterial haloalkane dehalogenase DhaA (EC 3.8.1.5) from *Rhodococcus rhodochrous* (Liss *et al*., 2015). The tag was modified to form covalent irreversible bonds with synthetic chloralkane ligands (Los *et al*., 2008; England *et al*., 2015). This covalent bond is formed rapidly under physiological environments and remains intact even under stringent conditions (Los and Wood, 2007). In addition, HaloTag^®^ proteins keep their monomeric structure, so the tag will not lead to oligomerisation of protein fusion partners (Banaz *et al*., 2018). Numerous ligands are available for the HaloTag^®^ including different fluorescent dyes with extended spectral range, photostability and membrane permeability such as the red fluorescent rhodamine derivative TMR (tetramethylrhodamine), the green fluorescent Oregon Green or the yellow fluorescent diacetyl derivative of fluorescein DiAcFAM. Furthermore, dyes can be varied in their dosages to either label all or only a few molecules which is needed for single-molecule tracking approaches. Reck-Peterson and colleagues (2006) for example used HaloTag^®^ and a TMR ligand successfully to label dynein in sea urchin axonemes and tracked single molecules with a precision of a few nanometres to reveal dynein’s stepping behaviour at microtubules. Moreover, several organic fluorophores especially for live-cell labelling and subsequent imaging were recently developed with photoactivatable properties (Lee *et al*., 2010) as well as with improved quantum efficiency and superior brightness while retaining excellent cell permeability (Grimm *et al*., 2015).

In 2012, Ishikawa and colleagues identified several split points within the HaloTag^®^ protein and demonstrated its reconstitution ability. These results gave rise to the idea of establishing the Split-HaloTag^®^ imaging assay *in planta* as the usability of the HaloTag^®^ imaging system in plants has been shown previously (Lang *et al*., 2006). This new Split-HaloTag^®^ approach is particularly useful for characterising the assembly protein complexes at structural elements including cell membranes or the cytoskeleton. The possibility of using these microscopy techniques will enable to study local formation of a given complex with improved details compared to BiFC and conventional confocal laser scanning microscopy. Single-molecule tracking approaches with low concentrated fluorescent dye will enable the tracing of complex mobility at the cytoskeleton or inside the membrane system. In this study, to establish the new Split-HaloTag^®^ imaging assay we used the previously described anchoring of the molybdenum cofactor biosynthesis complex via molybdenum insertase Cnx1 to filamentous actin (Kaufholdt *et al*., 2017). In this way, we demonstrate the advantages of this assay for imaging of *in vivo* protein-protein interactions via advanced microscopy.

## Material and Methods

### Cloning of Split-HaloTag^®^ Gateway destination vectors

The optimised HaloTag^®^-7 sequence (298 amino acids) from Promega (https://www.promega.de/) was genetically split on position 155/156 aa into the N-terminal fragment “NHalo” (aa 1-155) and the C-terminal fragment “CHalo” (aa 156-298) according to the initial experiment of Ischikawa and colleagues (2012). In order to create Split-HaloTag^®^ GATEWAY^®^ destination vectors, enabling C-terminal Split-HaloTag^®^ reporter fusion, the binary destination vectors pDest-*Cluc-*GW and pDest-GW*-Cluc* were used (Gehl *et al*., 2011). PCR-Primers were designed to fuse specific restriction enzyme recognition sequences at both reporter fragments (Table S1). After amplification, *Cluc* fragments were exchanged by restriction and ligation for *Nhalo* or *Chalo* residues using the restriction sites *Xba*I and *Spe*I (N-terminal) and *Xho*I/*Sac*I (C-terminal), respectively (restriction enzymes purchased by Thermo Fischer Scientific (https://www.thermofisher.com), to create pDest-*Nhalo*-GW, pDest-*GW*-*Nhalo*, pDest-*Chalo*-GW and pDest-*GW-Chalo* (Table S2).

### Expression vectors

Coding sequences of Cnx6 (AT2G43760), Cnx7 (AT4G10100) and Map65 (amino acids 340– 587; AT5G55230) were fused to Split-HaloTag^®^ fragments via a two-step fusion PCR with Phusion-Polymerase purchased from Thermo Fischer Scientific (https://www.thermofisher.com). For the first PCR, each single cDNA and reporter fragment was created with an overlapping sequence to each, which enable assembly of fusion constructs (used primers listed in Table S1). For the second step, the products of the first PCR were assembled due to the overlapping matching sequences and then amplified a one single fragment. This *att*B-site flanked constructs were subcloned via BP-reaction into the Donor vector pDONR/Zeo to create entry vectors. Recombining these into pK7WG2 (Karimi *et al*., 2002) using LR-reactions generated the expression vectors pExp-*Nhalo*-*cnx7*, pExp-*Chalo-cnx6* and pExp-*Chalo*-*map65*.

All BiFC expression vectors and entry vectors with coding sequences of Cnx1 (AT5G20990), LA (Lifeact; amino acids 1–17 of the *Saccharomyces cerevisiae* protein ABP140), ABD2 (aminoacids 325–687; AT4G26700), CKL6 (amino acids 302–479; AT4G28540) and NLuc were available and are described by Kaufholdt *et al*. (2016a). The entry vectors were used to clone Split-HaloTag^®^ expression vectors via LR-reactions into pDest-*GW*-*Nhalo* and pDest-*GW-Chalo* to create pExp-*cnx1*-*Nhalo*, pExp-*la*-*Nhalo*, pExp-*ckl6*-*Nhalo*, pExp-*la-Chalo*, pExp-*abd2-Chalo* and pExp-*Nluc-Chalo*.

### Plant transformation

*N. benthamiana* wild type plants were cultivated in soil under greenhouse conditions. They were used for *Agrobacterium-*mediated transient transformation of fusion constructs 7 to 12 weeks after germination as described by Gehl and colleagues (2011). *Agrobacterium* strain C58C1/pMP90 carrying binary expression vectors were freshly grown (48 h at 28 °C) on solid CPY media (0.1% (w/v) yeast extract, 0.5% (w/v) casein peptone, 0.5% (w/v) sucrose, 2 mg/L MgSO_4_ × 7H_2_O (pH 7); 1.5% w/v agar) containing rifampicin (50 mg/L) and gentamycin (50 mg/L) as well as kanamycin (50 mg/L). Helper strain p19 (Voinnet *et al*., 2003) was grown on CPY medium containing rifampicin (50 mg/L) and kanamycin (50 mg/L). After growing for 20 h in 9 mL of liquid CPY at 200 rpm at 28 °C, cells transferred into fresh activation medium (10 mM MES/KOH (pH 5.6), 10 mM MgCl_2_,150 µM acetosyringone). Before infiltration of the bacteria into the leaves, each strain was diluted in activation media to an optical density of OD_600_ = 0.9 (final OD_600_ = 0.3). Then, three strains were mixed for each transformation: (i) One strain containing a NHalo construct, (ii) one strain containing a CHalo-construct and (iii) the helper strain p19. After incubation for 2 h at 50 rpm (28 °C), mixed *Agrobacterium* suspension were infiltrated into abaxial site of young but fully expanded leaves. Plants were incubated for 3-5 days in the green house.

### Staining of *N. benthamiana* leaves discs with HaloTag^®^ ligands

Staining protocol was based on the work of Lang and colleagues (2006). Leaf discs of (6-10 mm) of *N. benthamiana* leaves were transferred into a 20 mL syringe with screw lid and infiltrated with 2-4 mL ligand solution (0.5, 1.0 or 2.0 µM TMR, DiAcFAM and Oregon Green in 10 mM MES/KOH (pH 5.6) and 10 mM MgCl_2_). All dyes were purchased from Promega (https://www.promega.de/). Syringes with leaf discs were wrapped in aluminium foil and incubated for 0.5, 15, 30 or 60 min either on the work bench, on a tumbling shaker or on a rotary tube mixer. After staining, the samples were washed with 10 mL washing solution by closing the screw lid and moving the plunger up and down for 10 times. Washing steps were repeated with fresh washing solution 6-12 times. Furthermore, one duration before the last washing step of 0, 3 or 12 hours in washing solution was performed.

### Confocal laser scanning microscopy

The confocal Laser Scanning Microscope LSM 510Meta from Zeiss (Göttingen, Germany) was used. The cLSM-510META scanhead was connected to the Axiovert 200M. All images were examined using either the Plan-Neofluar 10x/0.3 or the C-Apochromat 40x/1.2 water-immersion objective. For excitation, both an argon laser (488 nm for BiFC, Oregon Green and DiAcFAM as well as chlorophyll fluorescence) and a Helium-Neon Laser (543 nm line for TMR) was used. The emitted light passed the primary beam-splitting mirror UV/488/543/633 and was separated by a secondary beam splitter at 545 nm. Fluorescence was detected with filter sets as follows: BP 505-530 nm for BiFC (Em_max_: 515 nm), Oregon Green (Em_max_: 520 nm) and DiAcFAM (Em_max_: 521 nm); BP 560-615 for TMR (Em_max_: 578 nm); LP 650 nm for chlorophyll fluorescence. Bright field images were taken with the transmitted light photomultiplier. All images were taken using ZEISS Microscope Software ZEN 2009 and processed with ZEN lite and Fiji (Schindelin *et al*., 2012). The shown images depict represented cells of several analysed leaves from at least three independent transformations.

### Super-resolution by polarisation demodulation microscopy

Principles of the experimental set-up are described by Hafi and colleagues (2014) and modifications enabling the analysis of the dye’s 3D-orientations were described by Albrecht *et al*. (2020). The cover slip was fixed to the microscope slide with nail polish. Linearly polarised light deriving from a 488 nm continuous wave (CW) laser (sapphire 488-50, Coherent) was used for excitation of Oregon Green molecules. The beam was expanded through a telescope system. The polarisation was modulated at 15 frames per modulation period by rotation of a λ/2-waveplate. The rotation was achieved through a chopper wheel (Optical Chopper Systems, Thorlabs) which was synchronised to an electron-multiplying charge-coupled device (EMCCD) camera (iXonEM+897 back illuminated, Andor Technology). Through the rotation of 2 wedge prisms lateral shift of the beam was caused that enabled measurements of the fluorophores being excited from a different direction. Then, the beam was focused onto the back aperture of the microscope objective (UPlanSApo, 60×, NA = 1.35 oil immersion, Olympus), which was integrated in an inverted microscope body (IX 71, Olympus). Emitted light was then passed through a dichroic mirror (beam splitter z 488 RDC, AHF) and an emission filter (ET band pass 525/50, AHF). To further magnify the image and focus it the EMCCD camera an additional lenses system was used. During the measurement 2,000 frames at approximately 32 ms per frame were recorded. The first 200 frames were neglected for calibration purposes. The raw fluorescence intensity of all modulation periods of the last 1,800 frames of a measurement was used for analyses. Images were “deblurred” by using deconvolution algorithms. The “blurring” function or point-spread function (PSF) was approximated by using the PSF-Generator Plugin for ImageJ (http://bigwww.epfl.ch/algorithms/psfgenerator/). Using the PSF, the modulating fluorescence intensities were deblurred using an iterative least-squares deconvolution while accounting for the polarisation modulation. The least-squares functional was minimised by using the FISTA Algorithm (Beck and Teboulle, 2009).

## Results and Discussion

The Split-HaloTag^®^ constructs created in this study are based on the enhanced HaloTag^®^-7 sequence, which has been optimised and improved with regard to solubility, stability, binding kinetics and access to an optional TEV-cleavage site (Ohana *et al*., 2009). According to the initial experiment of Ischikawa and colleagues (2012) HaloTag^®^ protein was split on position 155/156 aa into the N-terminal fragment “NHalo” (aa 1-155) and the C-terminal fragment “CHalo” (aa 156-298). The stable bond between a HaloTag^®^ protein and its ligand is formed by the catalytic amino acid Asp^106^ as part of the “NHalo” fragment. In the wild type dehalogenase, the closely located His^272^ would catalyse hydrolysis of the intermediate, resulting in product release and enzyme regeneration (Los *et al*., 2008). In the mutated HaloTag^®^ protein, however, the substituted Asn^272^ (HaloTag^®^-7) as part of “CHalo” traps the reaction intermediate as a stable covalent adduct. Taking human embryonic kidney cells as a model system, Ishikawa and colleagues (2012) affirmed the general capacity of Split-HaloTag^®^ reconstitution by using a self-associating Split-GFP system. Furthermore, they monitored membrane fusion as cell fusion enabled functional HaloTag^®^ reconstitution resulting in TMR signals after staining. In this method paper, we set out to (I) examine the utility of the Split-HaloTag^®^ system as new tool to study protein-protein interactions in plant cells, (II) to develop staining and washing protocols resulting in low background fluorescence and (III) to demonstrate application examples of Split-HaloTag^®^ for sophisticated imaging and advanced microscopy methods.

### Reconstitution of Split-HaloTag^®^ *in planta*

The cDNAs of the interaction partners were fused N-or C-terminally to the NHalo and CHalo fragments via fusion PCR and gateway cloning (see Materials and Methods). For high flexibility, Split-HaloTag^®^ GATEWAY compatible destination vectors were generated which enabled a fast and easy cloning of expression vectors with coding sequences of different proteins of interest. The complex formation of the heterotetrameric molybdopterin synthase (MPT) subunits Cnx6 and Cnx7 from *Arabidopsis thaliana* was used to demonstrate the capability of reconstitution of the Split-HaloTag^®^ *in planta*. This protein pair was chosen as positive control due to their verified high binding strength (Kaufholdt *et al*., 2013). The two interacting proteins (Fig. 1A) will bring the two HaloTag^®^ reporter fragments in close spatial proximity and, thereby, guide the reconstitution of functional HaloTag^®^ proteins.

**Fig. 1:**
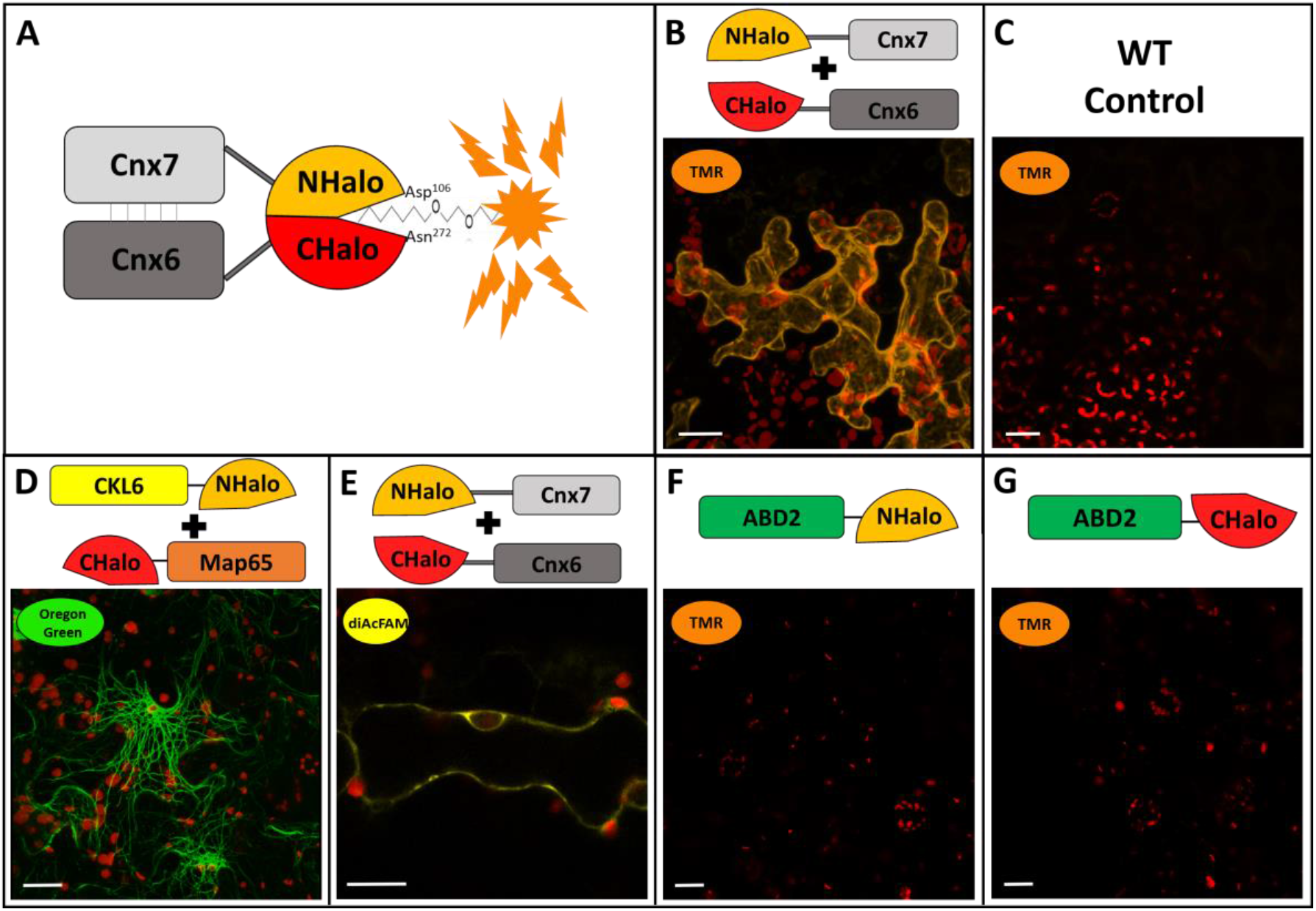
Testing Split-HaloTag^®^ complementation upon protein interactions of MPT synthase complex with different fluorescent ligands. Shown are images of *N. benthamiana* epidermis cells via confocal microscopy. Staining of leaf discs was performed 4-5 days after transformation. All images were taken with a C-Apochromat 40x/1.2 water immersion objective. Scale bars depict a length of 20 µm each. **(A)** Schematic illustration of Split-HaloTag^®^ reconstitution guided by the MPT synthase subunits Cnx6 and Cnx7. The important amino Asp106 and Asn272acids of each reporter terminus for covalent linker binding are depicted. **(B)** Cytosolic TMR fluorescence after transformation with NHalo-Cnx7 and Cnx6-CHalo. **(C)** Negative control of a wildtype (WT) leaf after staining. **(D)** Oregon green fluorescence at microtubules filaments after transformation of microtubules binding constructs CKL6-NHalo and CHalo-Map65. **(E)** Cytosolic diAcFAM fluorescence after transformation with NHalo-Cnx7 and Cnx6-CHalo. **(F/G)** Negative controls of single transformed Split-HaloTag^®^ reporter constructs fused to ABD2.

After transformation of *Nicotiana benthamiana* epidermis cells and staining with the fluorescent ligand TMR, specific cytosolic fluorescence was observed as a thin layer at the cell periphery (Fig. 1B). Wild type leaves were stained similarly, and lacking fluorescence signals indicated the washing steps being sufficient to remove unbound ligands (Fig. 1C). Therefore, since the MPT synthase complex is localised in the cytoplasm, HaloTag^®^ fragments CHalo and NHalo were capable of reconstitution guided by the strong interaction of Cnx6 and Cnx7. The reconstituted HaloTag^®^ was able to bind red fluorescent ligand TMR (Em_max_ 578 nm), green fluorescent ligand Oregon Green (Em_max_ 520 nm) (Fig. 1D) as well as yellow fluorescent ligand DiAcFAM (Em_max_ 526 nm) (Fig. 1E), even though DiAcFAM signals were weaker compared to TMR and Oregon Green when stained with similar concentration. This demonstrates a general binding capability of the reconstituted HaloTag^®^ exemplary for a plurality of other available dyes depending on the individual setting. As the two amino acids (Asp^106^ as part of NHalo and Asn^272^ as part of CHalo) important for covalent ligand binding are located on each of the two separate Split-HaloTag^®^ fragments, individual expression of “NHalo” or “CHalo” fragments will not enable ligand binding without each other which is a fundamental aspect when using Split-HaloTag^®^ imaging for investigation of protein-protein interactions (Fig. 1F/G).

After proving the general reconstitution with a protein pair forming a permanent complex, this new assay was tested in a second approach with MPT synthase subunit Cnx6 and molybdenum insertase Cnx1. In contrast to the permanent interactions within the MPT-synthase complex (Cnx6/Cnx7), interaction strength of the protein pair Cnx1/Cnx6 was previously found to be distinct but of a more transient nature (Kaufholdt *et al*., 2013). Like in previous BiFC and Split-Luciferase experiments (Kaufholdt *et al*., 2013), we again conducted a full PPI study including all necessary controls. As control proteins, the cytosolic proteins NLuc (N-terminus of the luciferase from *Photinus pyralis*) and the G-box protein GF14 (AT1G78300) were provided and both represent proteins showing no interaction with Cnx1 or Cnx6. To ensure staining with equal TMR concentration and a similar washing procedure leaf discs of interaction approach and controls were punched in slightly different sizes and stained simultaneously in the same syringe. The interaction approach showed strong cytosolic fluorescence (Fig. S1A) supporting Split-HaloTag^®^ reconstitution ability also for transient interactions. The negative control and both abundance controls showed weak cytosolic TMR signals, too (Fig.S1B-D). A negative control without any fluorescence would be an unrealistic event and observed spontaneous self-assembly was expected as it is typical for split-protein assays such as BiFC and Split-Luciferase when proteins are overexpressed in the small cytosolic space of plant cells (Gehl *et al*., 2009/2011). To evaluate whether differences in interaction approach and negative control are due to different protein concentrations we previously introduced additional abundance controls for our BiFC and Split-Luciferase studies (Kaufholdt *et al*., 2016b). The amounts of expressed proteins were therefore similar in all approaches. Taken all together, it can be concluded that random self-assembly can successfully be distinguished from real interactions. In comparison, BiFC fragments have an intrinsic affinity towards each other and once the BiFC complex is formed, this formation is irreversible (Kerppola, 2008). Irreversibility can also be assumed for Split-HaloTag^®^ since ligands will only covalently bind to reconstituted HaloTag^®^ proteins. However, with adequate negative controls and a careful evaluation of obtained results, the spontaneous self-assembly of both BiFC and Split-HaloTag^®^ experiments can be overcome. Using the transient interaction of the protein pair Cnx1/Cnx6 as an example we could validate the Split-HaloTag^®^ as a new addition to the large toolbox for investigation of PPIs *in planta*.

### Insights into the Staining Protocol

Lang and colleagues (2006), who first introduced the HaloTag^®^ system to plant cells, attributed great importance to washing procedures to reduce unspecific background fluorescence as background-less staining is often more complicated in plants than in animals. For interaction studies, comparison of fluorescence intensity and fluorescence pattern is the main task, and each form of background will falsify the result. However, after using the published destaining protocol an excess of unbound dye remained in the tissue. Therefore, optimised staining and destaining procedures had to be established for using Split-HaloTag^®^ as reliable tool for PPI studies. To improve staining protocols and to evaluate the best HaloTag^®^ ligand several different factors were investigated: (i) the size of analysed leaf discs (6; 8; 10 & 12 cm in diameter), (ii) concentration of ligands (0.25, 0.5, 1 & 2 µM), (iii) ligand incubation time (1 min up to 1 h), (iv) number of subsequent washing steps (6 up to 12), (v) incubation time in washing solution (3 h up to 12 h) and (vi) aeration of leaf discs in washing solution.

During this optimisation process, several observations were made that are worth mentioning and need to be considered to prevent misinterpretation of gained results. After TMR staining, fluorescence was always detected in vascular tissue of transformed as well as wild type leaves suggesting a nonspecific adhesion between TMR and molecules in leaf veins (Fig. S2A). Therefore, leaf area further away from vascular tissue should be used for analysis. Furthermore, staining with more than 0.5 µM TMR combined with an insufficient number of washing steps resulted in oversaturation and accumulation of unbound dye in the cytoplasm of parenchyma cells (Fig. S2B). This amount of unbound TMR accumulation increased when using larger or damaged leaf discs or older plants. Moreover, recycling of frozen TMR solution led to unspecific aggregations inside the cells.

DiAcFAM staining gained an overall weaker fluorescence signal compared to TMR but no staining of vascular tissue was observed (not shown). However, weak DiAcFAM signals (Em_max_ 521 nm) could easily be mistaken for typical plant background fluorescence at approx. 530 nm. Furthermore, accumulation of unbound ligands occurred especially in stomata after DiAcFAM (Fig. S2C) but also after Oregon Green (Fig. S2D) staining. In addition, Oregon Green resulted in accumulation inside vacuoles of parenchyma cells if incubated more than a few seconds in staining solution (Fig. S2E).

After testing of the different staining parameters optimal results for TMR staining were obtained using leaf discs of a diameter of 6-8 mm, a final TMR concentration of 0.5 µM in 2 mL fresh staining solution, 15 min incubation time and followed by eight subsequent washing steps into a 20 mL syringe and an overnight incubation in washing solution. Samples were incubated with 10 mL washing solution by closing the screw lid and moving the plunger up and down for approx. 10 times. Immediately before microscope analysis two more washing steps were performed. Both, overnight incubations either on a tumbling shaker or on a rotary tube mixer were equally sufficient for all dyes, as long as there was sufficient air in the syringe to allow leaf disc aeration. It must be noted that application of strong pressure during staining and washing procedure can cause severe stress and damage to the cells. Therefore, pressure to leaf discs in the syringe should be as low as possible but just enough for successful removal of all unbound ligand.

Oregon Green showed optimal results if a 0.5 µM staining solution was exchanged with 2 mL washing solution immediately after its infiltration and incubated for 15 minutes. Then, the washing procedure was applied as described for TMR. This could reduce but not completely avoid Oregon Green accumulation.

Overall, all exemplary tested dyes are suitable for intracellular cytosolic labelling of HaloTag^®^ proteins in plant tissue, which leads to the assumption that other HaloTag^®^ ligands will be suitable for other experimental settings as well. In this study, both TMR and Oregon Green showed ideal properties for confocal laser scanning microscopy despite Oregon Green accumulations. Due to a different experimental set-up only Oregon Green was used in SPoD microscopy. TMR showed the best applicability for usage within a Split-HaloTag^®^ complementation assay for staining *N. benthamiana* leaf discs.

### Validation of Split-HaloTag^®^ imaging upon protein-protein interaction at cytoskeletal elements

To further prove the usability of Split-HaloTag^®^ for *in planta* PPI studies, a third approach was conducted with proteins attached to cytoskeleton structures such as filamentous (F-) actin as well as microtubules. By applying this approach, it was possible to test whether the Split-HaloTag^®^ system allows for investigating the assembly of different proteins at cytoskeletal elements. Both structures were not labelled directly to NHalo or CHalo termini, but *via* binding proteins, as a fusion of larger reporter fragments directly to globular actin or tubulin proteins might disturb their polymerisation processes. For F-actin labelling, the binding domain of fimbrin from *A. thaliana* (ABD2; Sheahan *et al*., 2004) as well as of Abp140 from *Saccharomyces cerevisiae* (Lifeact/LA; Riedl *et al*., 2008) were used. Furthermore, microtubule binding domains of the two proteins Casein-Kinase-1-Like-6 (CKL6; Ben-Nissan *et al*., 2008) and the Microtubule Associated Protein 65 (Map65; Hamada, 2007) from *A. thaliana* were investigated. The cytoskeleton binding proteins show no direct protein interactions to each other. However, their affinity and subsequent binding and anchoring to the cytoskeletal structures results in such spatial proximity that it is able to act as model for a direct interaction of a cytoskeleton associated protein complex (Kaufholdt *et al*., 2016a). Expression of F-actin binding protein constructs followed by HaloTag^®^ reconstitution and staining resulted in a TMR specific fluorescence visible as transversely arranged filaments with branches distributed throughout the cytoplasm (Fig. 2A1). The approach with CLK6-NHalo and CHalo-Map65 to label microtubules upon Split-HaloTag^®^ reconstitution (Fig. 1D/Fig. 2A2) displayed filamentous structures more equally distributed throughout the cell with less cross bridges compared to actin filaments. These structures are typical for F-actin and microtubules, respectively, and were observed in BiFC experiments before (Kaufholdt *et al*., 2016a). In all interaction approaches both actin filaments as well as microtubules, could successfully be visualised upon interaction of cytoskeletal binding proteins.

**Fig. 2:**
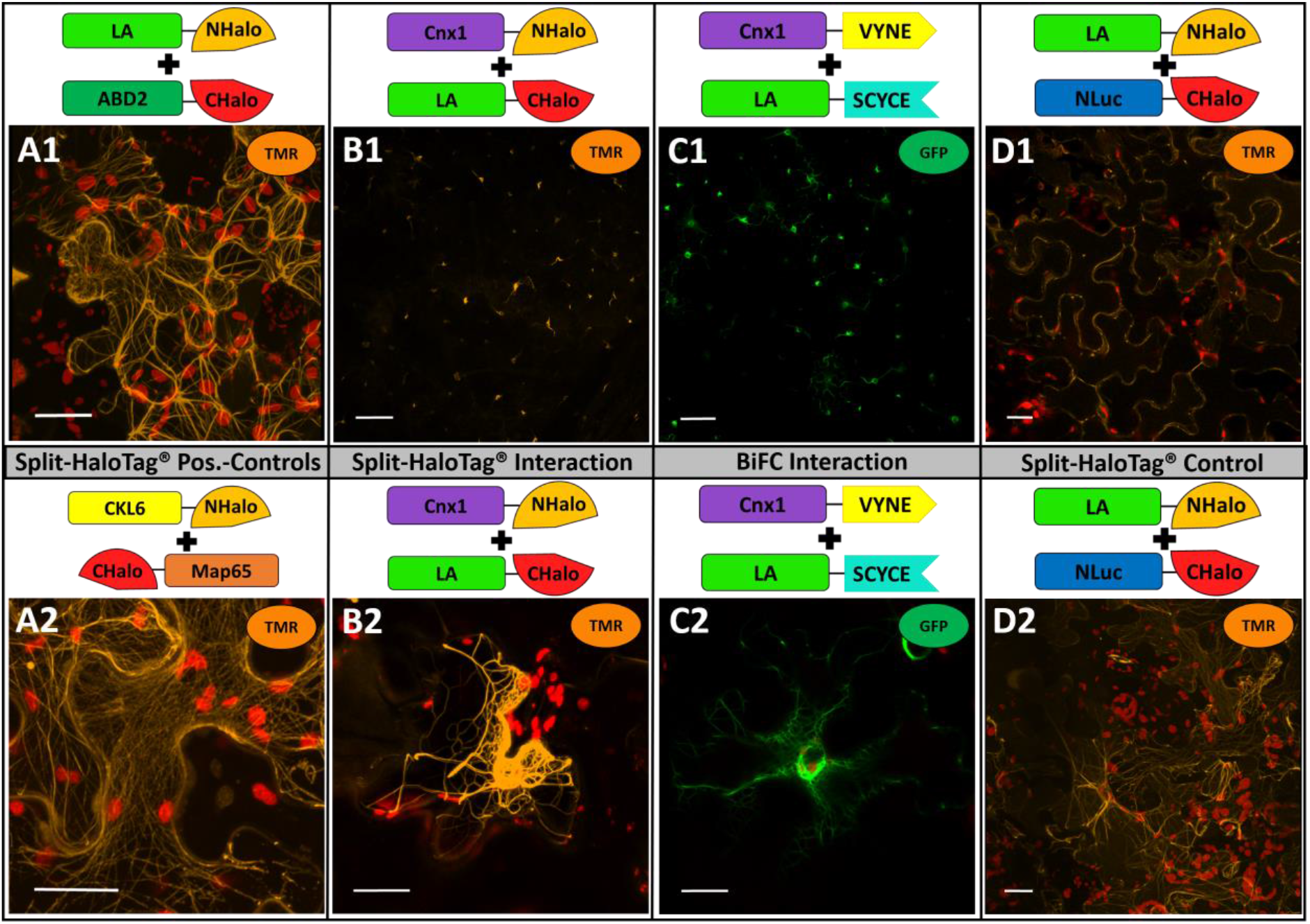
Split-HaloTag^®^ Protein-Protein Interaction Studies of Cnx1 and Actin Filaments via Lifeact. Shown are images of *N. benthamiana* epidermis cells via confocal microscopy of TMR or GFP. Staining of leaf discs was performed 4-5 days after transformation. All images were taken with a Plan-Neofluar 10x/0.3 **(B1/C1/D1)** or with a C-Apochromat 40x/1.2 water immersion objective **(A1/A2/B2/C2/D2)**. Scale bars depict a length of 100 µm **(B1/C1/D1)** or 20 µm **(A1/A2/B2/C2/D2). (A1)** TMR fluorescence at actin after transformation of actin binding constructs LA-NHalo and ABD2-CHalo. **(A2)** TMR fluorescence at microtubules filaments after transformation of microtubules binding constructs CKL6-NHalo and CHalo-Map65. **(B)** Split-HaloTag^®^ approach with Cnx1-NHalo and LA-CHalo. **(C)** BiFC approach with Cnx1-VYNE and LA-SCYCE. **(D)** Corresponding Split-HaloTag^®^ negative control where Cnx1 was replaced by the independent protein NLuc. **(B)** and **(C)** were imaged with identical setting for optimal comparison of strength and pattern.

Then, we aimed to validate this new assay with a binding assay of the molybdenum insertase Cnx1 to F-actin as a well-characterised example for a protein-protein interaction study at cell structures (Kaufholdt *et al*. 2016a). This established setting via cytoskeleton binding proteins for *in vivo* interaction studies was used to investigate whether the Split-HaloTag^®^ system demonstrate the F-actin binding of Cnx1 in the same manner it was shown by BiFC experiments before (Kaufholdt *et al*. 2016a). In the interaction approach, reporter fusion constructs of Cnx1 and the actin binding protein LA, respectively, were co-expressed in *N. benthamiana*. For comparison, the BiFC approach described by Kaufholdt *et al*. (2016a) was included, for which the reporter halves VYNE (N-terminus of Venus) and SCYCE (C-terminus of SCFP) were used (Gehl *et al*., 2009). Both BiFC and Split-HaloTag^®^ complementation assay show almost identical results (Fig. 2B/C). Both TMR and GFP fluorescence were detected in a filamentous pattern concentrated at the actin nucleus basket and thinned out at F-actin towards the cellular cortex. A typical pattern for studying an actin interacting protein complex was observed in both approaches that is reminiscent of a “starry sky” (Kaufholdt *et al*., 2016a) caused by F-actin anchoring of the interacting proteins in close proximity to its synthesis by the two actin binding domains of LA and Cxn1. When LA-NHalo was co-expressed with NLuc-CHalo in the negative control, TMR specific fluorescence at actin filaments was detected, too. However, TMR fluorescence was equally distributed within the cell and no starry sky could be observed (Fig. 2D1/D2). Consequently, identical results of both BiFC and Split-HaloTag^®^ were observed showing the characteristic “starry sky” like pattern demonstrating its interaction with F-actin as it was discussed before by Kaufholdt *et al*. (2016a) This proves exemplary the applicability of the Split-HaloTag^®^ system on the basis of a protein binding to the cytoskeleton.

### Application examples of Split-HaloTag^®^ for sophisticated imaging

The Split-HaloTag^®^ imaging assay has been proven as a feasible method for imaging of protein interactions and obtained results of all given examples demonstrate similarity to BiFC results. However, regarding protocol simplicity and handling, Split-HaloTag^®^ imaging assay cannot outcompete BiFC as method of choice for studying putative protein interaction. However, BiFC is limited when confirmed specific protein interactions need to be observed and imaged with greater detail. As conventional light and fluorescence microscopy are diffraction-limited, advanced fluorescence imaging methods such as single molecule detection, subdiffractional polarisation imaging or super-resolution microscopy (SRM) techniques have been developed to improve the resolution and to allow studying molecular processes more detailed (Moerner and Kador 1989; Orrit and Bernard, 1990; Bode *et al*., 2008; Holleboom *et al*., 2014; Liao *et al*., 2010/2011; Godin *et al*., 2014; Loison *et al*., 2018; Camacho *et al*., 2019). For such imaging techniques, stable fluorescent dyes emitting light at various wavelength are needed which is hard to realise by standard fluorescent proteins (Banaz *et al*., 2018). When BiFC would be used to localise and image a specific protein interaction, detected fluorescence intensity would be directly interlinked with fusion construct expression levels. Compared to this, the Split-HaloTag^®^ system has the advantage of adjusting the dosage of fluorescent dyes customised for the individual application. Using 0.25 µM compared to 0.5 µM for example enabled us to observe the attachment of molybdenum insertase Cnx1 to F-actin with much greater detail (Fig. 3A/B). Using even lower concentrations would label even less HaloTag^®^ proteins and enable tracking the dynamic movements of single protein complexes.

**Fig. 3:**
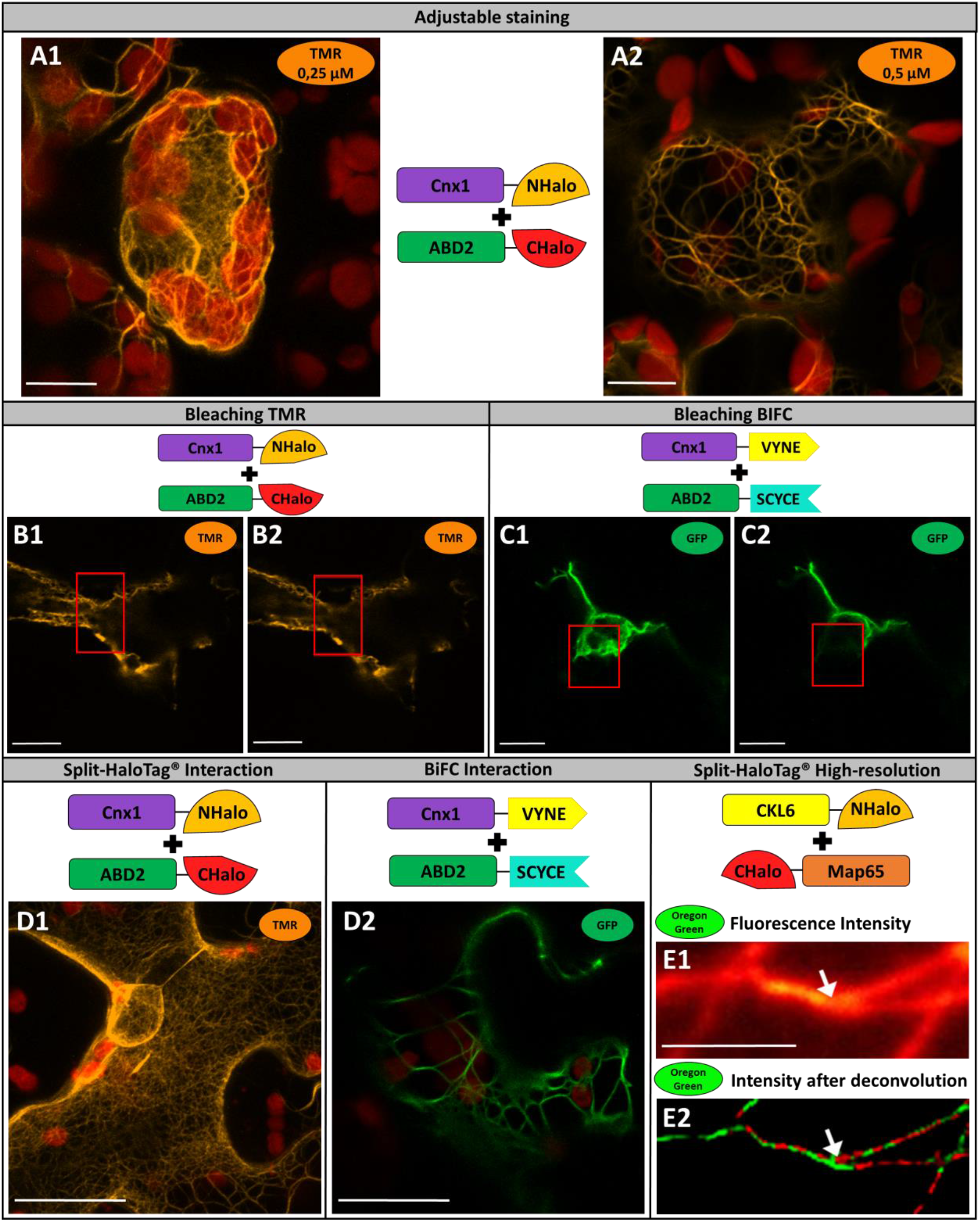
Analysis of Split-HaloTag^®^ images via confocal microscopy and super-resolution SPoD microscopy. Shown are representative *N. benthamiana* cells 4-6 days after transformation. **(A)** Staining with different concentrations of TMR (**(A1)** 0,25 µM and **(A2)** 0,5 µM) to optimise fluorescence intensity. **(B/C)** Bleaching experiments of **(B)** TMR and **(C)** BiFC. Shown are pictures before **(B1/C1)** and after **(B2/C2**) of 100 iterations of 100% laser power in the marked section (red rectangle). **(D)** Interaction studies at actin filaments with Cnx1 and ABD2 via Split-HaloTag^®^ **(D1)** or BiFC **(D2)**. Split-HaloTag^®^ Staining was performed with TMR. Images were taken with a C-Apochromat 40x/1.2 water immersion objective. Scale bars depict a length of 20 µm. **(E)** SPoD microscopy of microtubules stained with Oregon Green after transformation of the Split-HaloTag^®^ microtubules binding constructs CKL6-NHalo and CHalo-Map65. **(E1)** Diffraction limited image depicting averaged raw fluorescence intensity. **(E2)** Phase colour coded fluorescence intensity image after 1000 iterations of the deconvolution algorithm. The red/green colourcode support subdiffractional separation of the fibres at a distinct branching fork (arrow) that is not visible in the conventional diffraction of wide field image. However, future work is needed to enhance separation by different phases in addition to pure image deconvolution for entire cell images. The raw data scale bars depict 2 µm each.

Especially during long-term observations, stability of fluorescent dyes is of great importance. Even though FPs have improved a lot in recent years with regard to their photon budget (Kubitscheck *et al*., 2000) standard FPs used for most BiFC experiments show low quantum efficiency, blinking behaviour and a high photobleaching rate (Reck-Petersen *et al*., 2006). In a direct comparison, the GFP of the BiFC approach bleached much faster after 100 iteration (100% laser power) compared to TMR with same settings (Fig. 3B/C).

To show the stability and resolution potential of TMR for confocal laser scanning microscopy the interaction of Cnx1 and the actin binding protein ABD2 was again used as example. For this purpose, each layer of a cell needs to be scanned in very thin optical slices (µm range or less) and such a detailed imaging can take several minutes. The stable TMR fluorescence of a Split-HaloTag^®^ enabled more defined results of cell images (Fig. 3D1) compared to BiFC approaches (Fig. 3D2), which is demonstrated by the very thin filaments of the F-actin network hardly detectable by BiFC.

Super-resolution by polarisation demodulation (SPoD) microscopy was used as second example for demonstrating the performance of the Split-HaloTag^®^ system. This advanced fluorescence imaging technique is a subdiffractional polarisation imaging method that allows measurement of the average orientation of fluorescent dyes attached to different structures and was first described by Hafi and colleagues (2014). Fluorescent molecules are illuminated via linearly polarised light. This causes the fluorophores to be excited at different times, which results in a modulated fluorescence intensity from the fluorophores. Depending on the orientation of the illuminated fluorophores (or more specifically the orientation of their transition dipole moments), the observed fluorescence intensity will be phase-shifted, and differently oriented fluorophores will emit periodic signals peaking at different points in time. Therefore, the analyses via deconvolution algorithms allow a high-resolution imaging of cell structures (for details see Albrecht *et al*., 2020). During each measurement, 2,000 frames were recorded, which in itself is not problematic when using stable dyes. Split-HaloTag^®^ constructs with the two microtubule binding domains of CLK6 and Map65 were used for this purpose. Overexpression of MAP65 isotypes is known to result in microtubule bundling (Mao *et al*. 2006). After transformation and expression, leaf discs were stained with Oregon Green, which wavelength was more appropriate for SPoD microscopy. The observed fluorescence resulted in individual microtubules (Fig. 2E1). Albeit high amounts of background fluorescence and out-of-focus signal complicated the recording and modulation analysis, subdiffractional separation in a branching region of distinct fibres was observed in the deconvolved image (Fig. 3E2) and was supported by different phases as visualised by a simple red/green colour code. These different phases in the branching region of the two fibres were already observable in the raw modulation data. Certainly, future work is needed to enhance separation by different phases in addition to pure image deconvolution for entire cell images. Nevertheless, Split-HaloTag^®^ imaging assay can be used for such advanced fluorescence imaging techniques, which failed in case of BiFC.

### Conclusion

In this study, Split-HaloTag^®^ imaging assay was established for the first time *in planta*. Vectors were cloned and reporter termini NHalo and CHalo were tested for reconstitution in both fusion orientations to the protein of interest and in all four orientation combinations to each other. The applicability of the system for protein-protein interaction studies was demonstrated using previously published protein interactions forming the Molybdenum cofactor biosynthesis complex including its anchoring to F-actin. As TMR also penetrates peroxisomal membranes (Lang *et al*., 2006) as well as the nuclear envelope (unpublished data), interaction studies in other organelles would also be possible. Regarding protocol simplicity and handling, Split-HaloTag^®^ imaging assay cannot outcompete BiFC as method of choice for studying putative protein interactions. An additional infiltration of a fluorescent ligand into the cell with subsequent washing steps was a disadvantage compared to BiFC. However, relating to the background, Split-HaloTag^®^ shows the same performance as BiFC, as spontaneous self-assembly is typical for Split-protein assays when proteins are overexpressed in the small cytosolic space of plant cells. The benefit of the Split-HaloTag^®^ system lies in the ability to visualise confirmed specific protein interactions with advanced imaging techniques. Therefore, this system can be used in future for sophisticated imaging techniques such as 3D-microscopy, polarisation-microscopy, single-molecule tracking or super-resolution imaging methods that require brighter and more stable fluorescent markers. Localisation of protein complexes can be observed with the Split-HaloTag^®^ imaging assay in a distinct manner. In live-cell microscopy, the method combines *in vivo* split-reporter analyses with the previous shown advantages of the HaloTag^®^ like a large set of differently coloured fluorescent ligands, their photostability compared to fluorescent proteins and the ability to vary labelling intensity via adjusting the dosage of dyes independent from protein expression. In recent years, improved FPs have already been used for split-reporter applications (Xie *et al*., 2017), however, some drawbacks still remain. Therefore, this Split-HaloTag^®^ imaging assay provides a unique and sensitive approach for characterization of PPIs by combining all advantages given by the HaloTag^®^ system with the advantages of protein fragment complementation assays.

## Supplemental information

Supplemental data are available online.

Figure S1: Split-HaloTag^®^ Protein-Protein Interaction Studies of Cnx1 and Cnx6 analogue to the BiFC study in Kaufholdt *et al*. (2013).

Figure S2: Evaluation of HaloTag^®^ fluorescent Ligands TMR, DiAcFAM and Oregon Green.

Table S1: Primers for Cloning and sequencing of Split-HaloTag^®^ related constructs and vectors.

Table S2: Split-HaloTag^®^ Destination and Expression vectors used within this work.

## Supporting information

Supplemental_Split-Halo_Meinen et al

## Acknowledgement

We are grateful to Dr. Christin-Kirsty Baillie for scientific support and critical reading. We thank Tanja Linke for excellent technical work in our lab. This work was financially supported by the Deutsche Forschungsgemeinschaft (grant GRK2223/1) to RH and RRM.

## Author Contribution

RMM, HB, PJW, RRM, RH and DK planned and designed the project. RMM, JNW, MB, CT, SF, JS and DK cloned the vectors, performed plant transformation and the cLSM experiments. AA and RM handled SPoD microscopy. All authors analysed and discussed the results. RMM, JNW, RH and DK were primarily involved in drafting the manuscript and RMM and DK produced figures and tables. JS, HR, PJW and RRM critically read the manuscript and improved the text, all authors finalised it. RH and DK coordinated the work.

## Abbreviations

BiFC: bimolecular fluorescence complementation;
diAcFAM: diacetyl derivative of fluorescein;
FP: fluorescent protein;
SPoD: Super-resolution by polarisation demodulation;
TMR: Tetramethylrhodamine

