## Supplemental_Split-Halo_Meinen et al for "Split-HaloTag^®^ Imaging Assay for Sophisticated Microscopy of Protein-Protein Interactions *in planta*"

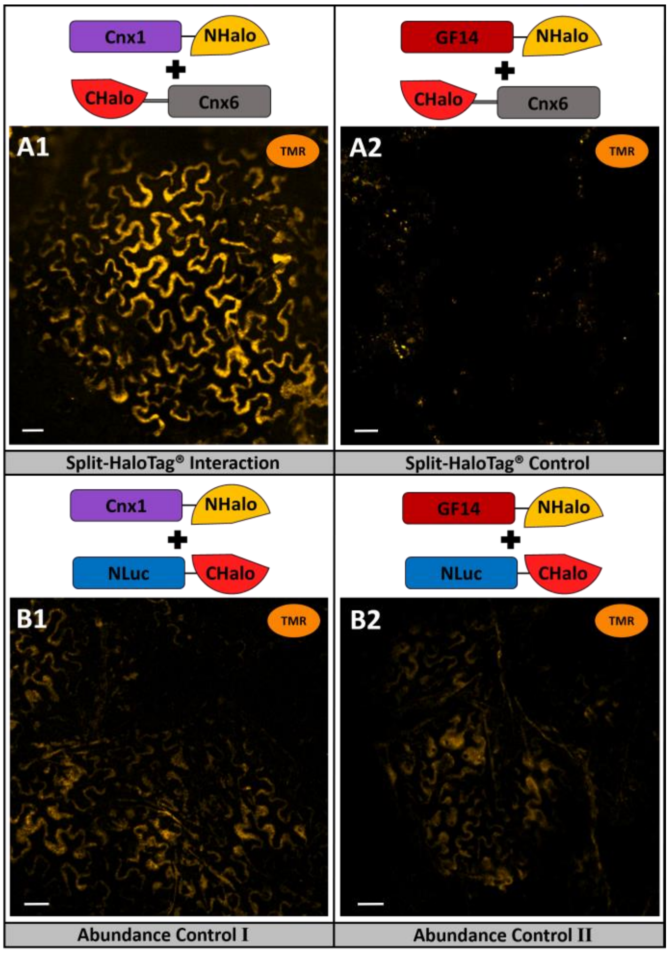


**Fig. S1: Split-HaloTag® Protein-Protein Interaction Studies of Cnx1 and Cnx6 analogue to the BiFC study in Kaufholdt *et al*. (2013).**

Shown are images of *N. benthamiana* epidermis cells via confocal microscopy of TMR. Staining of leaf discs was performed 5 days after transformation. All images were taken with identical setting (Plan-Neofluar 10x/0.3) for optimal comparison of fluorescence strength. Scale bars depict a length of 50 µm. **(A1)** Interaction approach with Cnx1-NHalo and CHalo-Cnx6. **(A2)** Negative control with the non-interacting proteins GF14-NHalo and CHalo-Cnx6. **(B)** Abundance control to validate negative control with **(B1)** Cnx1-NHalo and NLuc-CHalo as well as **(B2)** GF14-NHalo and NLuc-CHalo.

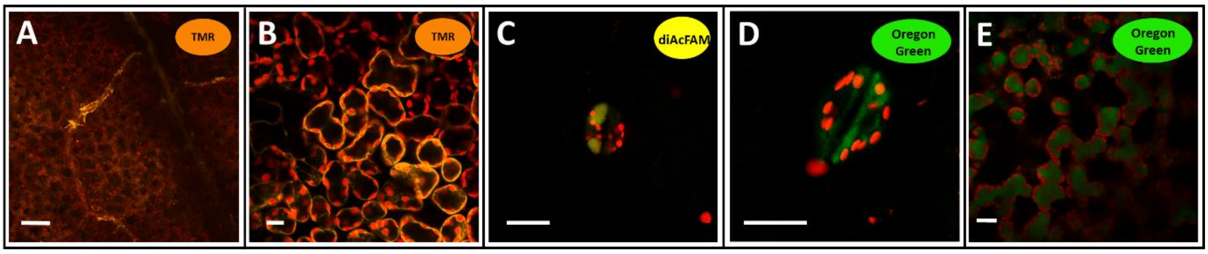


**Fig. S2: Evaluation of HaloTag® fluorescent Ligands TMR, DiAcFAM and Oregon Green.**

Confocal microscopy of *N. benthamiana* leaf discs stained with TMR **(A/B)**, DiAcFAM **(C)** or Oregon Green **(D/E)**. Images were taken either with a Plan-Neofluar 10x/0.3 **(A)** or with a C-Apochromat 40x/1.2 water immersion objective **(B-E)**. Scale bars depict a length of 100 µm **(A)** or 20 µm **(B-E)**. **(A)** Unspecific TMR binding in vascular tissue. **(B)** Oversaturation of unbound TMR in parenchyma cells after using 2 µM dye. **(C/D)** Accumulation of unbound ligands in stomata after DiAcFAM **(C)** and Oregon Green **(D)** staining. **(E)** Accumulation of unbound Oregon Green in parenchyma cells.

**Table S1: Primers for Cloning and sequencing of Split-HaloTag® related constructs and vectors.**
(**a**) Cloning of Split-HaloTag® Destination vectors (restriction + ligation); (**b**) Cloning of Split-HaloTag® Fusion Constructs. All oligonucleotides are purchased from Sigma-Aldrich (Steinheim, Germany).

| **Name** | **Sequence (5´-3´)** | **Purpose** |
|---|---|---|
| **XbaI_ATG_Nhalo_for** | GGGGGGTCTAGAATGGGATCCGAAATCGGTACTG | a |
| **NHalo_ Linker_SpeI_rev** | CAAAAACTAGTGCCGCTGCCGCCGGTGCGGAAGGCCTGGAAGGTC | a |
| **XbaI_ATG_Chalo_for** | GGGGGGTCTAGAATGACCGACGTCGGCCGCAAGCTGAT | a |
| **CHalo_Linker_ SpeI_rev** | CCAAAAACTAGTGCCGCTGCCGCCACCGGAAATCTCCAGAGTAGAC | a |
| **XhoI_Linker_NHalo_ for** | GGGGGGCTCGAGGGCAGCGGCGGCATGGGATCCGAAATCGGTACTG | a |
| **NHalo_N_SacI_rev** | CCAAAAGAGCTCTTAGGTGCGGAAGGCCTGGAAGGTC | a |
| **XhoI_Linker_Chalo_ for** | GGGGGGCTCGAGGGCAGCGGCGGCACCGACGTCGGCCGCAAGCTG | a |
| **CHalo-N_SacI_rev** | CCAAAAGAGCTCTTAACCGGAAATCTCCAGAGTAGACAGCC | a |
| **AttB1_N-Halo_for** | GGGGACAAGTTTGTACAAAAAAGCAGGTTTAACCATTGGATCCGAAATCGGTACTGG | b (*Nhalo-cnx7)* |
| **Cnx7Ueb_Linker_ N-Halo-rev** | CAATCTTTGTAACTTCTTTGTCCATGCCGCCGCTGCCGGTGCGGAAGGCCTGGAAGGT | b (*Nhalo-cnx7)* |
| **N-Halo_Linker_Cnx7Ueb_ for** | ACCTTCCAGGCCTTCCGCACCGGCAGCGGCGGCATGGACAAAGAAGTTACAAAGATTG | b (*Nhalo-cnx7)* |
| **Cnx7_Stopp_AttB2_ rev** | GGGGACCACTTTGTACAAGAAAGCTGGGTCTCAGCCGCCGCTTATCGGAGGTAT | b (*Nhalo-cnx7)* |
| **AttB1_C-Halo_for** | GGGGACAAGTTTGTACAAAAAAGCAGGCTTAACCATGACCGACGTCGGCCGCAAGCTGA | b (*Chalo-cnx6, Chalo-map65)* |
| **Cnx6Ueb_Linker_ C-Halo_rev** | GGTTCTTCTCCTCTGCAGACATGCCGCCGCTGCCACCGGAAATGTCCAGAGTAGACAGC | b *(CHalo-cnx6)* |
| **C-Halo_Linker_Cnx6Ueb_ for** | GCTGTCTACTCTGGAGATTTCCGGTGGCAGCGGCGGCATGTCTGCAGAGGAGGAGGACC | b (*CHalo-cnx6)* |
| **Cnx6_Stopp_attB2_rev** | GGGGACCACTTTGTACAAGAAAGCTGGGTCTCAAGAAGAAGATTTGTTATCTCCTGTAAT | b (*CHalo-cnx6)* |
| **Map65-3+4_Stopp_attB2_rev** | GGGGACCACTTTGTACAAGAAAGCTGGGTCTCATGGTGAAGCTGGAACTTGATG | b (*CHalo-map65*) |
| **Map65-3+4-Ueb_Link_C-Halo_rev** | CATGATTCTCTCACGAGCAGAGGCAGCGGCGGCACCGGAAATCTCCAGAGTAGACAGC | b (*CHalo-map65*) |
| **C-Halo-Ueb_Link_Map65-3+4_for** | GCTGTCTACTCTGGAGATTTCCGGTGCCGCCGCTGCCTCTGCTCGTGAGAGAATCATG | b (*CHalo-map65*) |

**Table S2**: **Split-HaloTag® Destination and Expression vectors used within this work.**

| Vector | Reporter | Fusion orientation | Selectable with |
|---|---|---|---|
| **pDest-*Nhalo-GW*** | HaloTag® (N-Terminus) | N-terminal | Kanamycin/Chloramphenicol |
| **pDest-*Chalo-GW*** | HaloTag® (C-Terminus) | N-terminal | Kanamycin/Chloramphenicol |
| **pDest-*GW-Nhalo*** | HaloTag® (N-Terminus) | C-terminal | Kanamycin/Chloramphenicol |
| **pDest-*GW-Chalo*** | HaloTag® (C-Terminus) | C-terminal | Kanamycin/Chloramphenicol |
| **pExp-*Nhalo-cnx7*** | HaloTag® (N-Terminus) | N-terminal | Spectinomycin |
| **pExp-*Chalo- cnx6*** | HaloTag® (C-Terminus) | N-terminal | Spectinomycin |
| **pExp-*Chalo-map65*** | HaloTag® (C-Terminus) | N-terminal | Spectinomycin |
| **pExp-*cnx1-Nhalo*** | HaloTag® (N-Terminus) | C-terminal | Kanamycin |
| **pExp-*la-Nhalo*** | HaloTag® (N-Terminus) | C-terminal | Kanamycin |
| **pExp-*ckl6-Nhalo*** | HaloTag® (N-Terminus) | C-terminal | Kanamycin |
| **pExp-*la-Chalo*** | HaloTag® (C-Terminus) | C-terminal | Kanamycin |
| **pExp-*abd2-Chalo*** | HaloTag® (C-Terminus) | C-terminal | Kanamycin |
| **pExp-*Nluc-Chalo*** | HaloTag® (C-Terminus) | C-terminal | Kanamycin |
| **pExp-*cnx1-vyne*** | Venus (N-Terminus) | C-terminal | Kanamycin |
| **pExp-*abd2-scyce*** | SCFP (C-Terminus) | C-terminal | Kanamycin |
| **pExp-*la-scyce*** | SCFP (C-Terminus) | C-terminal | Kanamycin |
